# Topological Structure of Population Activity in Mouse Visual Cortex Encodes Visual Scene Rotations

**DOI:** 10.1101/2023.02.13.528247

**Authors:** Kosio Beshkov, Gaute T. Einevoll

## Abstract

The primary visual cortex is one of the most well understood regions supporting the processing involved in sensory computation. Historically, our understanding of this part of the brain has been driven by describing the features to which individual neurons respond. An alternative approach, which is rapidly becoming a staple in neuroscience, is to study and analyze the geometry and topology of the manifold generated by the neural activity of large populations of neurons.

In this work, we introduce a rigorous quantification of the structure of such neural manifolds and address some of the problems the community has to face when conducting topological data analysis on neural data. We do this by analyzing publicly available two-photon optical recordings of primary mouse visual cortex in response to visual stimuli with a densely sampled rotation angle. Since the set of twodimensional rotations lives on a circle, one would hypothesize that they induce a circle-like manifold in neural activity. We confirm this hypothesis by discovering a circle-like neural manifold in the population activity of primary visual cortex. To achieve this, we applied a shortest-path (geodesic) approximation algorithm for computing the persistent homology groups of neural activity in response to visual stimuli. It is important to note that the manifold is highly curved and standard Euclidean approaches failed to recover the correct topology.

Furthermore, we identify subpopulations of neurons which generate both circular and non-circular representations of the rotated stimuli, with the circular representations being better for angle decoding. We found that some of these subpopulations, made up of orientationally selective neurons, wrap the original set of rotations on itself which implies that the visual cortex also represents rotations up to 180 degrees.

Given these results we propose that population activity can represent the angle of rotation of a visual scene, in analogy with how individual direction-selective neurons represent the angle of direction in local patches of the visual field. Finally, we discuss some of the obstacles to reliably retrieving the truthful topology generated by a neural population.

## 1 Introduction

In recent years, neuroscience has seen a surge in new recording techniques based on both electrophysiological [1, 2] and optical [3, 4] methods. While in the past it was only possible to record a few neurons at a time, the new techniques of today allow for the simultaneous recording of an immense number of neurons during behavior. This methodological change has also resulted in a conceptual shift away from interpreting the properties of single cells and rather to focusing on using new mathematical tools to analyze large groups of cells [5, 6]. Some examples of such tools are *representational similarity analysis* [7, 8], *machine learning models* [9, 10, 11, 12, 13] and *neural manifolds* [14, 15, 16, 17, 18, 19, 20] among others. While such tools are both fascinating and useful, the results are more difficult to interpret than single-cell recordings, and their verification can present many conceptual and methodological challenges.

Neural manifolds are of specific interest in this work. They are a tool in which neural activity is binned across time or a set of stimulus parameters. The response in each bin can then be interpreted as a point in a high-dimensional vector space in which the neurons form a basis, we will also refer to these points as *states*. This representation of the activity of neural populations allows one to treat neural activity as a manifold, whose structure is connected to the computation implemented by said population. An especially important type of structure to study is the topology [21], which reflects the way in which different states of activity of the neural population relate to each other [22, 23].

Studying the topology of neural manifolds has produced many insights. Some examples of this are: ring-like manifolds in the mouse head direction circuit [24], toroidal manifolds in rat medial entorhinal cortex [25, 26, 27] and topological organization of place cell maps [28, 29, 30, 31]. These examples are all related to navigation or movement, but similar success stories seem to be lacking when it comes to both sensory and higher-order cognitive capacities. While fascinating work on higher-order cognition is on-going [32], understanding it in topological terms is quite a daunting task. Instead we focus on arguably the most widely studied sensory modality - vision, the analysis of which has become widely popular in light of several publicly available large-scale recordings [33, 34, 35].

### 1.1 Previous work on the topology of neural manifolds in vision

There are several papers in which researchers have used dimensionality reduction to study neural manifolds generated in visual tasks [36, 37]. In the case of sinusoidal gratings, dimensionality reduction has been used to identify a circular manifold [38]. More advanced dimensionality reduction techniques have been used to decode a toroidal manifold in primary visual cortex induced by the periodic structure of phase and orientation of drifting gratings [39, 40]. Furthermore, there have been several theoretical works proposing different models for the topological and geometric structure of activity generated by the visual cortex [41, 42]. One other paper worth mentioning in this context is Guidolin et. al. [43], in which the authors try to understand the geometry of visual space by combining spiking metrics [44] with topological methods. Nevertheless, the reduction methods used in these examples rely on visual inspection and remove structural properties of the data, due to the fact that the data has to be projected to a low-dimensional subspace. We address this point further in the Methods (Section 4).

To our knowledge only two papers have tried to quantify the topology of primary visual cortex in response to visual stimulation. The first paper to do this is [45], where the authors found a combination of spheres and circles generated by population activity in the primary visual cortex of macaque in a spontaneous and a natural movie condition. This analysis, while interesting, was limited by the fact that only a subset of five cells were used to generate the points ending up in the final analysis. Furthermore, such a result is very difficult to interpret as there is no clear way to extract and conceptually relate the spontaneous and natural movie conditions to their corresponding network states.

The second paper to do this is [46], which made use of the publicly available Allen visual coding neuropixel dataset [34] to study the topology generated by drifting gratings in various hierarchically organized lower and higher mouse visual regions. In this case a much larger set of cells were sampled, and one would expect that the drifting gratings generate a circular manifold in neural activity. While circles were often observed, especially when the temporal frequency of the drifting gratings was fixed, the analysis also seemed to reveal many other much more complicated topological structures.

In our judgement, while both of these works provide an interesting direction for future research, they are still inconclusive as to the precise nature of the topology of neural manifolds generated in visual tasks, even in very simple contexts like the presentation of grating stimuli. Furthermore, both of these analyses make use of electrophysiological recordings, whereas optical imaging methods might be better suited for this type of analysis as they allow for the recording of many more cells.

### 1.2 Our contributions

The availability of openly accessible large-scale recordings of neural data is a truly watershed moment for the development of neuroscience, as it allows the field to transition from generating hypotheses about cognitive function based on single neurons or very small populations, to thinking and hypothesizing in terms of neural populations made up of thousands of cells. In the present paper we contribute to this development by showcasing the topological structure we identify in mouse primary visual cortex from the optical imaging dataset made openly available in [35]. Our results show that when using an appropriate metric one can recover the circular topology in neural population activity that is generated by rotating images, which is not the case when using the straight line (Euclidean) metric. By an appropriate metric we mean that the distances between any two points are given by the shortest path between them which does not leave the surface of the manifold (unlike the Euclidian metric where the straight line may cross this surface). See Methods (Section 4), for a more detailed description.

Following this, in Section 2.2 we show that subpopulations contain different representations of the presented stimuli and that the angle of a rotated stimulus can be recovered more reliably from subpopulations which generate a circular topology. One of these subpopulations is of particular interest as it contains neurons with high orientation selectivity, which generate a circular manifold that wraps around on itself and is homeomorphic (has the same topology) to the real projective plane in one dimension. This observation raises interesting questions regarding the topology of neural activity generated by more complicated transformations of three-dimensional objects.

In Section 2.3 we show that recovering the expected circular manifolds in mouse V1, requires that both the number of cells and the angles of the presented stimuli are sampled densely enough. Finally, in the discussion 3.1, we outline three obstacles to the identification of topology in the visual system. We propose some potential solutions to these obstacles, which involve precise prescriptions about the type of experiments which are needed to overcome them and push the field further.

## 2 Results

As mentioned previously, the following results involved the analysis of the publicly available dataset [35], the preprocessing of which is described in detail in the Methods (Section 4). In Figure 1, we attempt to explain the main concepts involved in describing the topology of a neural manifold with the help of persistent (co)homology, as well as the extension of the geodesic metric to this method. The fundamental idea is that, stimuli can be thought of as coming from a *stimulus manifold* 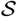 parametrized by some variables *θ*_1_,*θ*_2_, …,*θ_M_*. Then a population of neurons made up of *N* cells, responding to stimuli sampled from the *stimulus manifold*, traces out a surface in an *N*-dimensional vector space spanned by the firing rates of each neuron. In more precise terms a neural population applies a mapping Φ, which sends the values of the stimulus parameters to the space of possible population responses ℝ^*N*^, formally 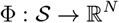. The surface traced out by this map Φ, is referred to as the *neural manifold*.

**Figure 1:**
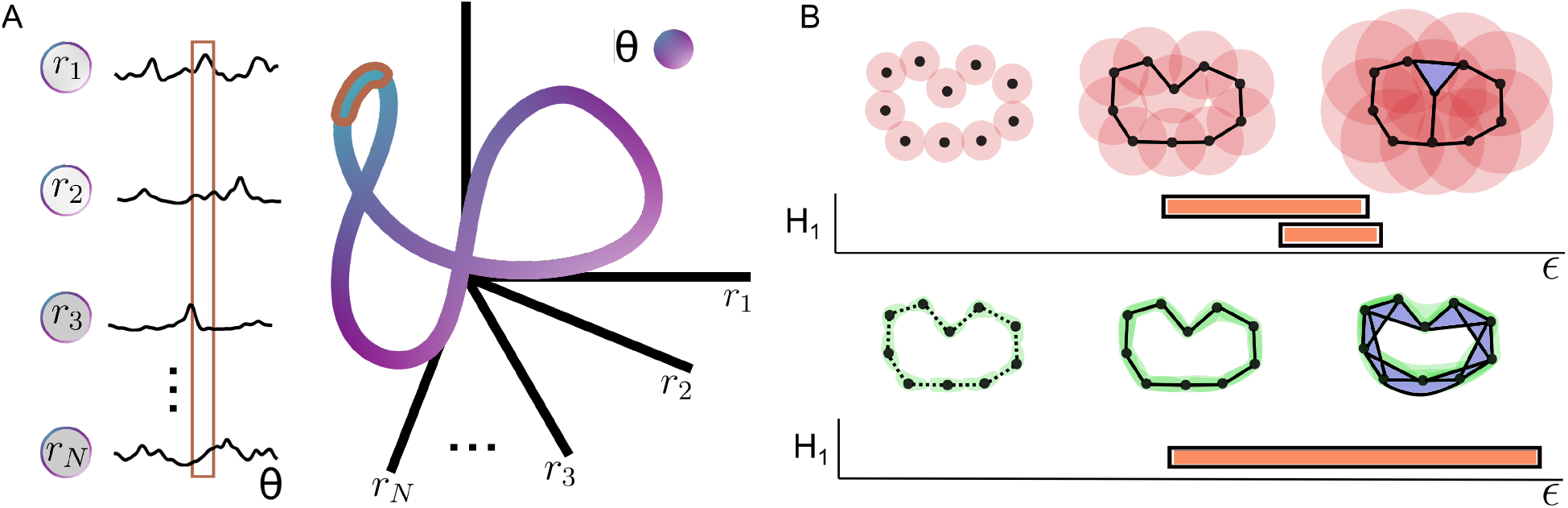
Neural manifolds and persistent (co)homology. A) Illustration of the concept of a *neural manifold*. The responses of a large number of neurons *N* to a stimulus parameter *θ* (**left**), can be thought of as a point in an N-dimensional vector space, spanned by the unit firing rate of each individual neuron. Taking all of these responses together, traces out a *neural manifold* (**right**). Similarly, the responses of a population to a range of stimulus parameters [*θ_t_, θ*_*t*+Δ*t*_], denoted by the brown box over the neural responses, gets mapped to a subset of the manifold, outlined by brown. B) An illustration of the persistent (co)homology algorithm along with a barcode, which is a visual representation of the algorithms’ output in which long bars correspond to highly persistent features. The red circles show how the Ripps complex is computed with the Euclidean metric. The green regions show the same for the geodesic metric, with the punctured edges corresponding to the graph generated by the initial choice of k-nearest neighbors (in this case k=2). Both the different metrics and the persistent (co)homology algorithms are described in further detail in the Methods (Section 4)

There are many ways to determine when two manifolds are different, but here we focus on their topology, or more precisely their (co)homology [47], which in simplified terms counts how many holes of different dimensions there are in the manifold. For example a frisbee, is a connected surface and therefore has no holes except one trivial zero-dimensional hole. A more formal way to express the same idea is to say that it has only one non-trivial zeroth (co)homology group (one way to write this with integer coefficients is 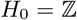 and *H_i_* = 0 ∀_*i*_ > 0), which is known to measure the number of connected components of a space. On the other hand a hula hoop, has two non-trivial (co)homology groups, namely the first and the zeroth (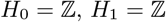 and *H_i_* = 0 ∀_*i*_ > 0). Therefore, it not only has one connected component, but also has a hole in its center. This concept of holes can be further generalized to any dimension, although the intuition of what a hole is fails to carry over for dimensions higher than two.

If one has a correct representation of a manifold, the (co)homology groups can usually be computed analytically. However neural data comes in the shape of a point cloud, which is why one has to input additional structure to the points. This is typically done with the help of persistent (co)homology, which is described in detail in the Methods (Section 4). The intuition behind this method is that by growing neighborhoods of size ε around each of the points (see panel B of Figure 1), one obtains a continuous surface in which the number of holes can be computed. Then the holes which last for a wide range of ε, are called persistent and correspond to a real topological feature of the data, unrelated to the inherent noisiness in the data generating process. This algorithm relies on the computation of a distance matrix, which is dependent on the choice of a metric.

A visualization of the experimental pipeline and the main result is shown in Figure 2. Mice (*N_m_* =3-6 depending on the stimulus condition), are presented with rotating stimuli while calcium imaging is used to record the population activity of tens of thousands of cells in their primary visual cortex (for more details see [35]). After this, responses were averaged within bins corresponding to ≈ 0.05 radians of the stimulus angle, shown in Figure 2C and described in detail in Methods (Section 4).

**Figure 2:**
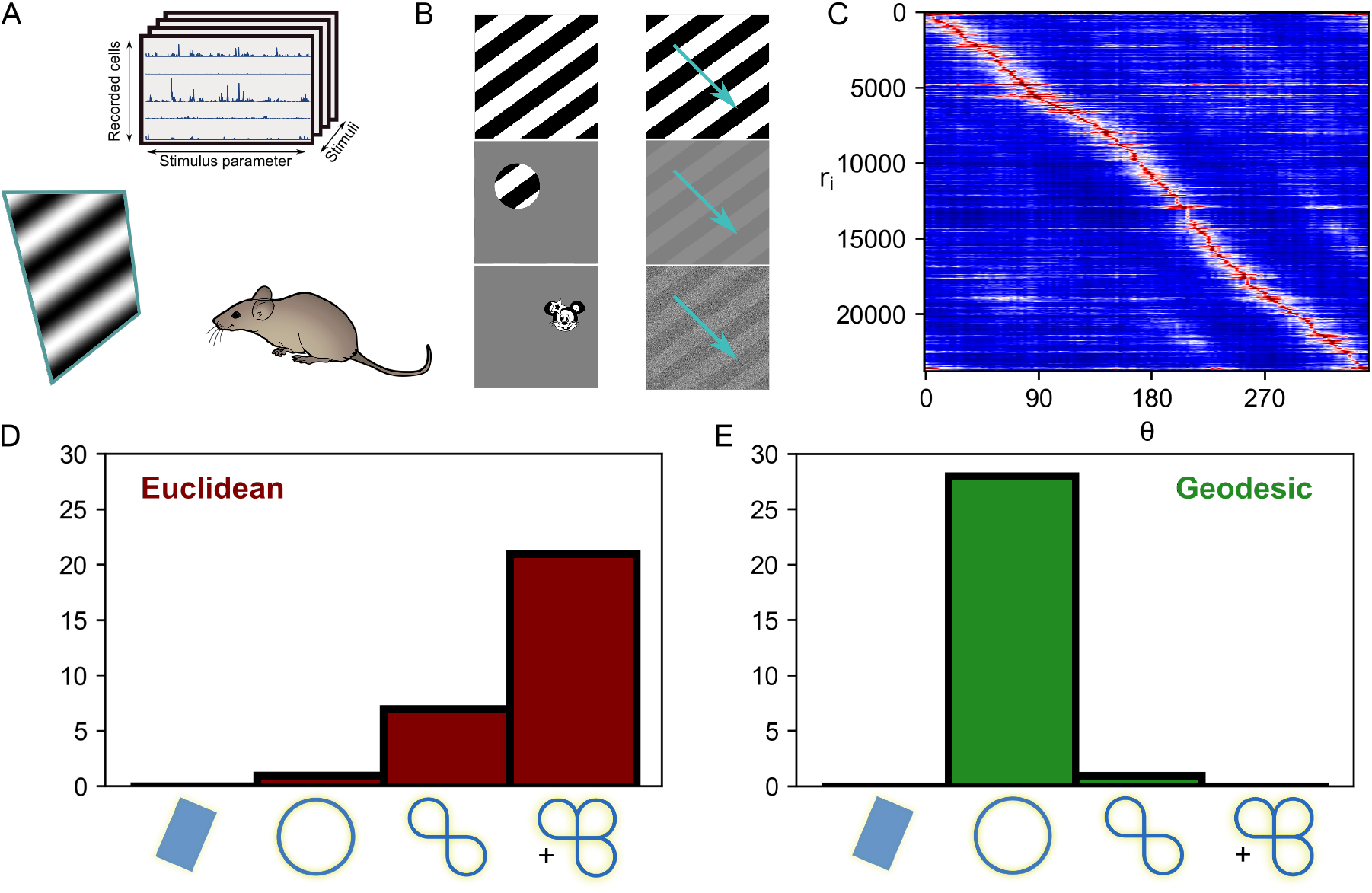
Data processing pipeline and significant (co)homological features. A) Illustration of the obtained data (**top**) and the experimental setup (**bottom)**. Stimuli are presented for 750 ms, along with 650 ms of gray screen between them. The response to each stimulus trial is defined as the summed activity during the first 650 ms after stimulus onset. See [35] for more details about the experiment. The mouse image was sourced from scidraw.io B) Example stimuli which were presented in the experiment. From left to right and top to bottom: static gratings, drifting gratings, local gratings, low contrast drifting gratings, image of Minnie and noisy drifting gratings. The stimulus set also included short static gratings and random phase sinusoidal gratings (not shown). All non-localized grating stimuli have a spatial frequency of 0.05 cycles per degree. The temporal frequency of the drifting gratings is 2 Hz. C) Neural responses binned by the angle of stimulus rotation and sorted by maximal response angle. The bin size for an angle was approximately 0.05 radians. D) Histogram of the identified manifolds across all stimuli and mice, when using the Euclidean metric. The last bin corresponds to cases in which the manifold has more than three holes, or in other words the first Betti number, defined in the Methods (Section 4), was larger or equal to three *β*_1_ ≥ 3. These results show that in most cases neural population activity lives on on a manifold with more than three holes. E) Histogram of the identified manifolds across all stimuli and mice, when using the geodesic metric.

### 2.1 Rotating stimuli generate circles in primary visual cortex

In order to identify the topology of the manifolds generated by neural activity, we used an adaptive k-nearest neighbor approximation of geodesics to compute a distance matrix, described in more detail in the methods (Section 4). Afterwards we computed the persistent (co)homology groups based on these distances. Histograms of the significant first (co)homology groups for all presented stimuli are shown in Figure 2E.

Our results, shown in Figure 2 panel E, point towards the fact that neural population activity in primary visual cortex responding to rotating stimuli lives on a circle in a high-dimensional vector space. This was true not only for symmetric grating-like stimuli, but also for non-symmetric localized ones.

Thereby one might think of primary visual cortex as encoding the angle of rotation of a visual scene. This result was observed in all stimulus conditions, except for one low-contrast stimulus presentation. In that case our statistical procedure determined that there are in fact two circles (see Figure 8 for the barcode plots).

Unlike the results above, which were obtained from using a geodesic metric, use of the Euclidean metric led to much more complex and varied results (see Figure 2 panel D) which means that to identify such structure one requires an approach which is invariant to the high degree of curvature generated by neural activity. The geodesic metric managed to deal with this curvature much more appropriately.

### 2.2 Subpopulations generate different representations of the circle

Unlike neural circuits involved in higher order cognitive capacities, the responses in the visual system are strongly stimulus driven [48]. This fact implies that the manifolds that one observes are less likely to be due to intrinsic attractor dynamics [49, 50] and more likely to reflect some transformation on the stimulus manifold. Given that in this dataset, all stimuli can be parametrized by a single angle parameter. It seems redundant that one needs on the order of 10^4^ neurons, to encode a one-dimensional manifold. However, a region such as primary visual cortex does not have to only encode the angle of rotation, but can be used to employ more complex transformations on the incoming inputs, while a smaller subpopulation encodes the relevant circular variables.

This line of thinking lead us to study subpopulations of neurons which exhibit peculiar tuning properties. To identify these subpopulations, we computed the orientation selectivity index (OSI) and the direction selectivity index (DSI) for each cell (see Methods in Section 4). Afterwards we identified four groups of cells, based on the most extreme values of these measures (|OSI_*s*_| > 0.3 and |DSI_*s*_| > 0.3, with 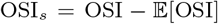 being a standardized selectivity index). One might argue that this way of choosing cells can bias the analysis, due to the fact that a very unbalanced number of cells can fall into each group. To account for this we also did the same analysis in which we picked the top 100 cells with the most extreme OSI and DSI values (see supplementary information Figure 7).

We call the first such group *direction-orientation (DO*) cells, in reference to the fact that they have both a high DSI (> 0.3) and a high OSI (> 0.3). We expected this group to reliably generate a single circular manifold. The second group, *orientation (O*) cells, have a large OSI (> 0.3) but a low DSI (< –0.3) and therefore a bimodal tuning curve. This is the most peculiar case, since if a cell responds in the same way to two rotations which are 180° apart, then the generated manifold should still be a circle but with opposite points identified. The third identified group had a high DSI (> 0.3) and a low OSI (< —0.3). We refer to cells in this group as *direction (D*) cells since while they show some tuning there is no easily extractable structure in their tuning curves. Finally, neurons with both a low OSI (< —0.3) and DSI (< —0.3), show no tuning and are therefore called *untuned*. Since the activity of these neurons are unrelated to the angle of the presented stimulus, it would be very unexpected for them to consistently encode circles.

As can be seen from Figure 3, *DO* and *O* cells predominantly generate circles, whereas the other two groups are more likely to generate topologically trivial manifolds (in other words spaces which can be contracted to a point, as they do not have any holes), with *D* cells rarely generating more complex topological structure. This is expected, as such cells fail to encode the circular structure of the stimulus manifold. To verify that *O* cells generate a circle which wraps around on itself, we repeated our analysis while limiting the presented angles to 180°. The histogram in Figure 3, panel D confirms this hypothesis. This can also be seen by the change in connectivity structure of the thresholded circular plots in the same figure.

**Figure 3:**
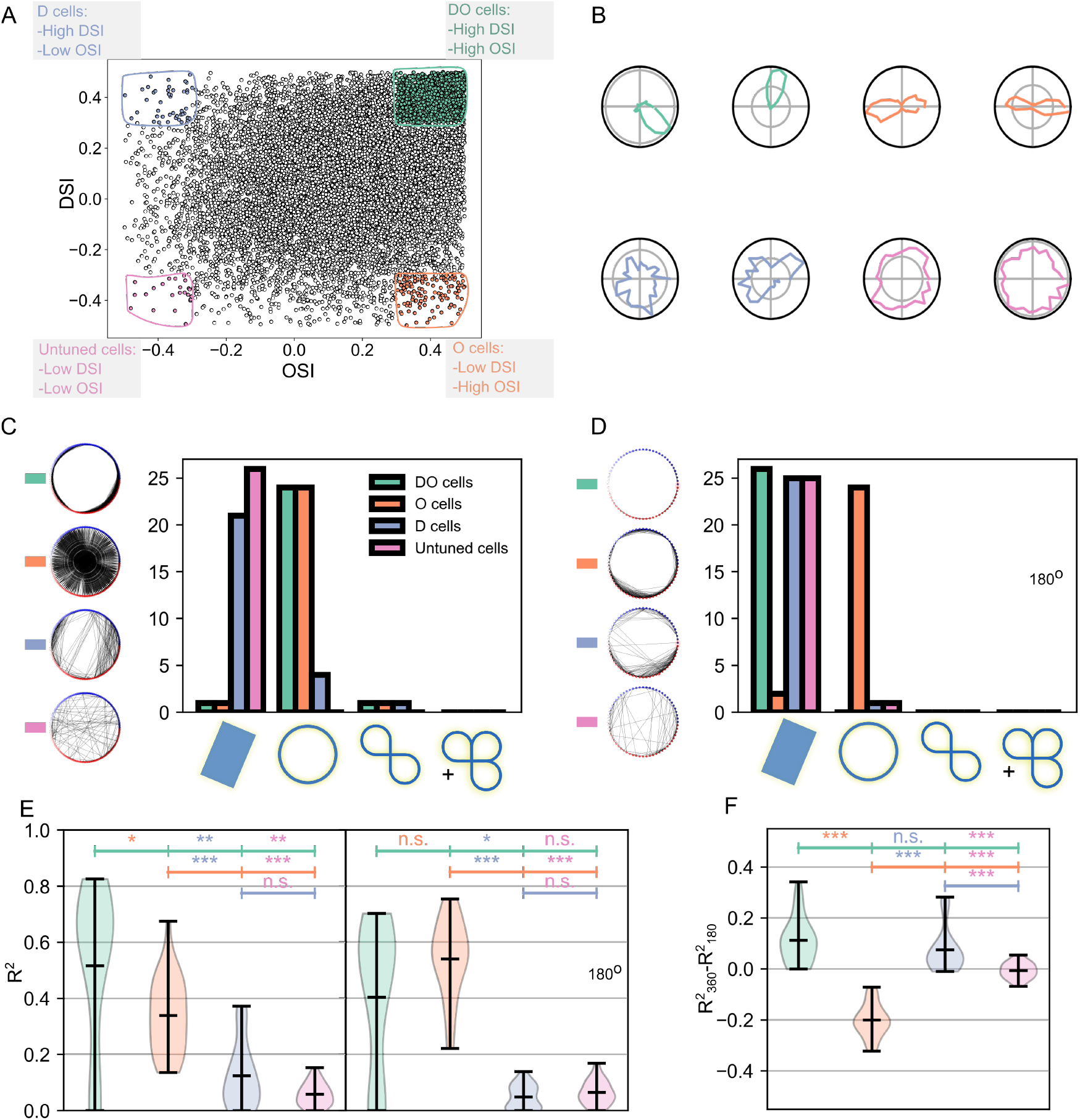
Subpopulations generate different manifolds. A) Scatter plot showing the focus cell groups, along with both the DSI and OSI of all cells. Each focus cell group is highlighted in a different color. B) Example polar tuning curves color coded for each cell group. C) **Left** - example circular graph plots showing the thresholded connectivity between points, in which points are organized in a circle and are connected by edges if they are closer than the chosen threshold. The threshold was chosen to be the birth plus one fourth of the persistence of the largest component of the first (co)homology group. **Right** - histogram of the identified manifolds for each cell group. D) Same as panel C, except the presented angles were limited to 180°. Furthermore, points at which the manifold fails to connect and complete the circle, are marked by red. E) Ordinary least squares decoding performance of the different subsets of cells. To determine significance we used a Bonferroni corrected Wilcoxon rank-sum test. *DO* and *O* cells showed good performance, whereas less strongly tuned cells under-performed. The left panel shows predictions for the full 360° of directions, whereas in the right panel the directions were recomputed as *θ* = *θ* mod 180°. F) The difference between the predictions showed that only O cells exhibit a significant improvement in their performance when it came to recomputed stimuli.

While projections generated by dimensionality reduction algorithms can be informative, they might also obscure important structural information even in the case of low-dimensional manifolds. Such methods have been widely critiqued [51, 52], and our analysis does not necessitate such a processing step as both distances and (co)homology can be calculated in the high-dimensional embedding space. Instead of performing dimensionality reduction and showing low-dimensional projections of the high-dimensional manifold, Figures 3 and 4 show circular graphs, in which an edge is drawn when the distance, computed in the high-dimensional embedding space, between two population states is lower than a chosen threshold.

**Figure 4:**
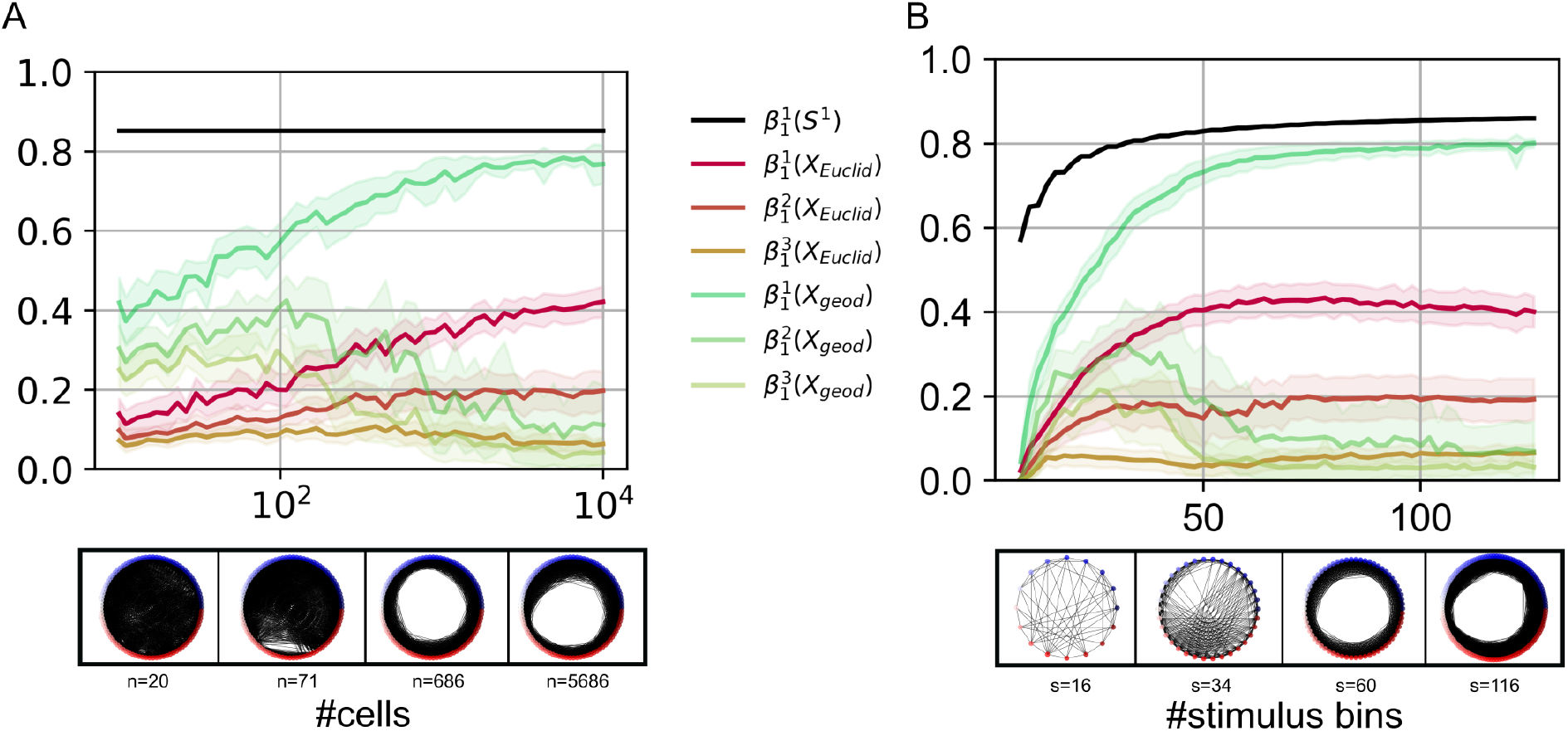
Sufficient sampling is necessary for the recovery of neural topology. A) **Top** - Normalized persistence of the top three one-dimensional Betti numbers/features, denoted by 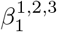, across all mice and stimuli responding to rotating stimuli whose angles were binned into 90 bins as a function of the number of subsampled cells. The black curve shows the persistence of the first one-dimensional Betti number of 90 points organized in a circle. Green curves correspond to the feature persistence after distances were computed using the geodesic metric, whereas the red curves were generated after using the Euclidan metric. **Bottom** - Plots of the thresholded connectivity between state vectors, with *n* being the number of neurons, representing different angles of a stimulus. The threshold was chosen to be half of the maximal distance between points. For all of these plots an approximation of the geodesic metric was used. B) **Top** - Normalized persistence of the top three features across all mice and cells responding to rotating stimuli as a function of the number of bins in which the stimuli were split. **Bottom** - Thresholded connectivity of the activity generated by the full population in response to a stimulus binned into different numbers of angles, with *s* being the number of binned angles.

Additionally we, use ordinary least squares (OLS) regression to show that the angle decoding performance, defined as the coefficient of determination *R*^2^ on a test set, described further in Methods (Section 4), of *O* cells improves significantly when the angles are set {*θ* → *θ* mod *π*}, in other words they obey the relation *θ* = *θ* + *π*, whereas this does not happen in the case of the other subpopulations. It is in stark contrast with both *DO* and *D* cells for which performance degrades as a result of this identification, leading to further support for the interpretation that *O* cells truly encode angles on a circle wrapped around on itself. In general, this analysis also supports the idea that cells participating in the generation of the circular manifold are responsible for encoding more information about the angle of rotation of a stimulus.

### 2.3 Identifying circles in V1 requires sufficient stimulus sampling

To study the amount of sampling necessary to reliably recover the topology generated by a neural population, we varied the bin size and subsampled different amounts of neurons. In Figure 4, we show how much sampling of both cells and stimuli is necessary in order to reliably recover the topology of the circular neural manifold. These results are shown for both the geodesic (green) and Euclidean (red) metrics. As a comparison baseline, we also plotted the persistence of an artificially created perfect circle (black). From these plots one can see that, with the geodesic metric, somewhere on the order of 1000 cells are necessary to reliably recover the topology of the circle, a manifold which is not particularly complicated. When it comes to the sampling of stimulus space, our results indicate that one needs to sample at the very least more than 60 angles to be confident in the estimated topology. Furthermore, in both cases using the Euclidean metric ascribes a large persistence value to not only one, but two one-dimensional features. This happens even when a high number of cells and stimuli is sampled.

It is worth noting that, while the aforementioned numbers are relevant for future experiments of this type, the general amount of sampling necessary to reliably extract the topology of a neural manifold might also depend on other factors. Examples of such factors are the curvature of the neural responses, the dimension of the stimulus space, the complexity of the task and others. We elaborate on some such factors in the Discussion (Section 3.1).

## 3 Discussion

In the present work, we have applied and extended tools from topological data analysis to large-scale neural recordings. We have shown that rotating visual stimuli consistently induce manifolds with a circular topology in primary mouse visual cortex. Furthermore, our analysis shows that subpopulations of cells classified based on traditional tuning-curve statistics, like the OSI and DSI, support the generation of circular, projective and flat manifolds in response to such images. This analysis was only possible due to the availability of large-scale neural recordings containing not only a dense sampling of both stimuli and cells, but also the choice of an appropriate metric, namely an approximation of the geodesic metric.

Following this, an important question for future research is what experiments should one perform to capture the full complexity of population activity during the performance of a task? It is crucial to note that, potentially with the exception of highly stable attractor networks, it is unclear how one can begin to disentangle the external input that a task induces in a population from the internal activity generated by said population. Therefore, the structure that one finds in a population performing a particular task can not be generalized to other tasks, except when an immense number of possible tasks are presented, which is incredibly difficult, if not practically impossible, to do in an experimental setting. Still one has to wonder whether there might be settings in which task spaces are particularly well suited to be sampled sufficiently densely to obtain an understanding of the system under study.

These are difficult questions and additional work will be needed to answer them fully. In the following discussion section we try to shine light on some of the obstacles inhibiting our ability to dissect all of this complexity, by understanding the conditions for which one can faithfully recover the correct topology of a neural manifold.

### 3.1 The sampling obstacle

Previous work by [53] has proposed the notion of *neural task complexity* (NTC) which bounds the dimensionality of neural activity. A consequence of this work is that besides the obvious dimensionality bound given by the number of recorded neurons, the complexity of the task which an animal is engaged in also plays a fundamental role in determining the properties of the final neural manifold which one can experimentally observe. Taking this point of view, one has to accept that when an experiment is done and neural activity is recorded, the manifold that is extracted is dependent, not only on the number of recorded cells, but also on the structure of the task.

Nevertheless, due to the particular structure of some tasks, the neural manifold generated by a population can be studied. For example, as mentioned before, in tasks related to motor action [15] and navigation [28, 24, 26, 25], very interesting structure has already been extracted. Such tasks have two significant advantages. The first is that they are inherently low-dimensional and as a result sufficiently sampling them is not as difficult. The second advantage is that in such tasks, a very large number of intermediate states are sampled. For example, in the case of head direction, being in a state characterized by a given angle requires that the animal goes through all intermediate angles. The same is true in the case of movement in a maze, where getting from point A to point B requires the representation of all intermediate states. In other words, due to the nature of these tasks, the stimulus space is always sampled densely.

These two advantages are not present in the case of the visual system, where the dimensionality of visual space is much higher. An estimate of the lower bound on the dimensionality of this space, or rather the space of natural scenes, is given by the dimensionality of Imagenet [54] which is 43 [55]. Although this is most likely a huge underestimate of the true dimensionality of visual space, it is nevertheless way too high to sample densely in an experimental setting. This dimensionality problem can often be avoided, by restricting the stimulus space to gratings, which can be parametrized by only a few parameters like orientation, phase and spatial frequency. However, even such simple experiments often fail to have the second advantage, as often only a few of these stimulus parameters are sampled. Therefore, both low-dimensional stimulus manifolds and dense sampling might be necessary.

These problems are not unique to vision, but can also appear in other sensory modalities and higher cognitive functions. Thus future work, trying to understand the neural manifolds generated by neural populations, should carefully consider both the structure and the density with which a task can be sampled and would also benefit from first studying stimulus spaces which can be densely sampled.

### 3.2 The metric obstacle

After population activity is recorded and a point cloud of population firing-rate vectors is generated, this collection of points is simply a set and there is yet no notion of how these points relate or how distant they are from each other. In order to talk about neural *manifolds*, such a notion is necessary (a more rigorous treatment of this is given in the supplementary information 6.1). At first glance, the most parsimonious assumption about these distance relationships is that they should obey the Euclidean metric. As elaborated in Section 4.4, this metric does not reflect the possible trajectories between two points, since the transformations which neural populations implement on the initial stimulus manifold are often highly nonlinear [56, 57, 58] and straight lines between points might correspond to impossible network states.

The most appropriate metric to use would be the geodesic metric [59], as it reflects the distance a state has to travel to get to a new state, but calculating it in even the simplest neural network models requires knowing all their parameters and solving a very high-dimensional differential equation. Doing this in actual data is not currently feasible. Nevertheless such a metric can be approximated using a *k*-nearest neighbor shortest path algorithm which we describe in the Methods (Section 4). A potentially more robust alternative to this approach, based on the UMAP algorithm, has also recently been proposed in [26].

The geodesic metric is more appropriate to use when one wants to know the topological and geometric properties of the neural manifold extracted in an experiment. However it is unclear whether the approximated geodesic metric can be used on the neural manifold generated in a different experimental setting. This is because in each experiment, the possible population states generated by a task do not exhaust all possible population states in general. Thus, different metrics can be used in tandem to answer different questions regarding *off-* and *on-manifold* perturbations. With the geodesic metric reflecting on-manifold perturbations and the Euclidean metric off-manifold perturbations. In this case by an off-manifold perturbation we mean the following.

#### Definition

Given an element *s* from a stimulus manifold 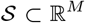 embedded in M-dimensions, an off-manifold perturbation is a perturbation *γ* such that 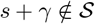.

In contrast, an on-manifold perturbation is a perturbation which does not end up outside the manifold.

While the Euclidean metric does not reflect the true distance between stimulus representations, when the stimulus space is restricted, this metric can still be used to describe how easy it would be to transform one stimulus into another using off-manifold perturbations. For example, in the data that we have analyzed, the stimulus manifold is given by the set of rotated images 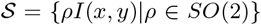, where *SO*(2) is the 2D rotation group containing all rotations in two dimensions and *I*(*x,y*) is an image. This means that any rotation of an image would be considered an on-manifold perturbation and the geodesic metric should be used to compare the populations responses to such perturbations. However one could also add noise, scale or even use generative models to interpolate between images in a complicated manner [60, 61, 62], which would all impose a different metric on the data. Since the Euclidean metric always uses the shortest paths between population states in the embedding space, it can also be used to describe the closest states in terms of the off-manifold perturbations between them.

Given this task dependence of the metric, future research could focus on generating tasks in which the stimulus manifold is given interesting topological structure and the change in the optimal metric is studied. Additionally, it might be worthwhile to use modelling and simulations of artificial neural networks in order to understand how network properties relate to the topological properties of the neural manifolds they generate.

#### 3.3 The decoding obstacle

Once a population generates a particular manifold, one might wonder how this encoding of stimulus information is processed by downstream neurons and is typically studied by training a single layer linear decoder [63]. Since different subpopulations within a region can apply different transformations to a stimulus manifold, there is always the possibility that by analyzing the responses generated by the whole recorded population one misses interesting topological structure in a subpopulation which is processed individually in downstream layers. In the present work we were able to rely on widely accepted measures of visual tunning curves, with which to identify the subpopulations. However these measures do not exist for arbitrary tasks and regions and may not be the most appropriate way to understand neural decoding. It is even possible that we have failed to identify more complex and interesting topological structure in the data.

To address such worries, in the future it will be highly beneficial to apply clustering algorithms which are able to extract topologically different and decoding-relevant subpopulations. An approach popular in topological data analysis, known as cohomological decoding [64, 65] has already been applied in the context of neuroscience [66, 26, 20] and future studies can further take advantage of such methods. In the context of grid cells, previous work has made use of clustering methods to identify cells whose activity plays a role in generating a torus [25] (the neural manifold regularly observed in grid cell recordings). Such cells have even been found to play a more fundamental role in encoding position in artificial recurrent neural network models performing a path integration task, than cells with a high grid-score [67].

#### 3.4 Outlook

Building a theory of neural population function, by studying how such populations transform stimulus manifolds into neural manifolds, with potentially different topological or geometric properties, is a young and fascinating research program. Despite the myriad of publications and findings about neural manifolds, pushing this program to more complex and scientifically revelatory settings, will not be a trivial task. With the hope of contributing to the development of this research program, we have gone into detail about some of the potential obstacles one might run into when trying to understand neural function in these terms. To address how these obstacles might be overcome, we propose that in order to reliably obtain the true topological structure of a neural manifold, future experiments have to prioritize sampling the stimulus manifold under study densely and choosing an appropriate metric for the analysis of the data. Furthermore, we assert that the field would strongly benefit from topologically inspired clustering algorithms, which will help reveal more fine grained structure in population activity.

## 4 Methods

### 4.1 Data and preprocessing

We obtained the data from Pachitariu et. al.[33]. The recording and preprocessing of this dataset is described in detail by the authors in [35]. Here we describe the additional processing we applied to study topological structure.

In order to obtain a topologically robust point cloud, we split the angles of each image into 126 bins, each one of size ≈ 0.05 radians and averaged all population responses in these bins. This left us with a 126X#neurons matrix for each image/animal condition, to which we could then apply persistent homology.

In order to extract the four subpopulations which ended up in our final analysis, we first computed the OSI and DSI as,

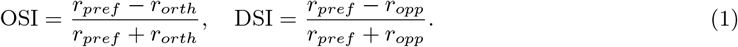

With *r_pref_, r_orth_, r_opp_* being the preferred, orthogonal and opposite orientations respectively. We further standardized these measures (OSI_*s*_ and DSI_*s*_) in the range [−0.5, 0.5]^2^ by subtracting their mean. For the analysis shown in Figure 3, we then only considered strongly selective cells in the union of subsets corresponding to different cell classes [−0.5, −0.3]^2^ ∪ ([−0.5, −0.3] × [0.3, 0.5]) ∪ ([0.3, 0.5] × [−0.5, −0.3]) ∪ [0.3, 0.5]^2^.

For the decoding analysis, described in the next section, one might argue that to fairly compare decoding performance of the different cell classes, each class should contain the same number of neurons. To verify whether this could make a difference, we sorted the neurons into four groups, depending on which quadrant they occupied in OSI_*s*_ × DSI_*s*_ coordinates. After this we sorted them by their *l*1 norm in this coordinate system.

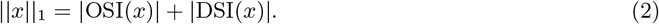

After this we selected the top 100 cells for further analysis (shown in SI Figure 7).

### 4.2 Dimensionality of the data

As mentioned in the introduction one might argue that when the set of visual stimuli is simple enough dimensionality reduction methods are sufficient to study the structure of neural activity directly by visualizing it, since only a few components are necessary to explain a large portion of the variance. However visual responses to simple stimuli, while intrinsically low dimensional, seem to posses a high degree of curvature and as a result it is not clear that such preprocessing steps are always justified.

In the presently analyzed data we z-scored each binned dataset and applied principal component analysis (PCA) to it. We observed that the first 2 components managed to explain 36% ± 8% of the variance on average, whereas using the top 10 components explained 79% ± 5%. This supported the idea that one should prefer a dimensionality-free topological approach.

One might still object, that while linear methods might not be appropriate, nonlinear techniques like Isomap [68], t-SNE [69] and UMAP [70] should do the job. For one, these methods are much harder to understand and verify and that is enough of a reason to avoid them when less agressive alternatives are on the table. However even ignoring that, they failed to visually reproduce the correct manifold structure by generating more than one circle on several datasets. Some visual examples are given in supplementary Figure 5. It is worth noting that these failures are quite rare especially when using Isomap.

Instead, for the purpose of visualization, we opted to use thresholded circular graphs, in which one creates a graph by considering each point in a point cloud as a vertex and drawing an edge between two points whenever the true distance between them in the high-dimensional space is lower than a specified threshold. Afterwards the graph is embedded in two dimensions and organized in a circle, which is especially convenient when one is exploring circular manifolds. Given this organization, if a circle is present then the edges only connect neighboring vertices and do not cross over to the other side. The presence of edges crossing over hints that the manifold is not circular. The big advantage of this approach is that one can directly see the connectivity in circles in which opposite points are identified, which happens to be the case in *O* cells and can be directly seen in Figure 3 panel C. The threshold was chosen to be one fourth of the persistence of the biggest circular features plus its birth, which is when the circle appears clearly while avoiding unnecessary clutter. This visualization procedure was done using the Networkx Python package [71].

### 4.3 Linear decoding analysis

In order to compare the decoding performance across different subpopulations we randomly split all response/angle pairs into a train (80%) and a test (20%) set. Then we performed linear regression in order to predict the angle of each stimulus from the subpopulation response.

To evaluate the performance of the linear decoder, we evaluated the coefficient of determination *R*^2^, given by the formula,

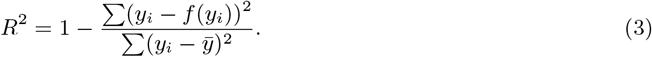

Where *x_i_* and *y_i_* are the input and target points to a statistical model applying the map *f* (·).

To determine significance we used a Bonferroni corrected, Wilcoxon rank-sum [72] test on each pair of subpopulation *R*^2^ distributions.

While non-linear decoders can surely improve the decoding performance, they are harder to justify biologically and add additional assumptions [63], which are outside of the scope of this work.

### 4.4 Computing metrics on neural manifolds

Metrics, also known as distance functions, are a way in which to compute the distance between two points in any space. The most famous such metric is the flat/Euclidean metric, which is simply given by the expression,

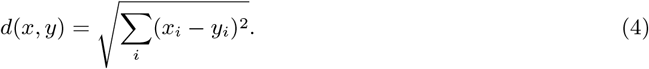

This is a good metric when neural activity lives in flat space, but in the case of a more complicated manifold with curvature, a geodesic metric is more appropriate. This type of distance is given by the expression,

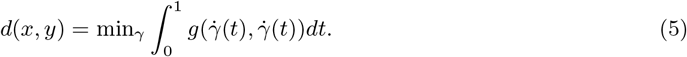

Where *γ*(*t*) is any curve connecting *x* and *y* on the manifold and *g* is the metric tensor, for more information see [59]. While this is the “correct” metric, it is only possible to obtain an expression for it if one knows the precise expression which generates neural activity and even then solving this optimization problem is non-trivial. In the case of neural recordings, such an expression definitely does not exist, therefore we opt for a numeric approach with which to approximate these geodesics. This approach has previously been applied in [46], with the slight difference that the nearest neighbor graph was constructed by fixing a distance threshold instead of finding the k-nearest neighbors, and is similar to newer developments in which the UMAP algorithm has been adapted to approximate geodesics [26].

The geodesics are approximated by the following stepwise algorithm:

1. Given a point cloud *X* embedded in an n-dimensional space ℝ^*n*^, choose a number of neighbors *k* and construct the *k*-nearest neighbors graph.
2. Compute the distance between each pair of points by calculating the shortest path on the graph between them.
3. (*adaptive - optional*) If *k* was chosen too low, there might be disconnected clusters of points between which the distances are infinite. To fix this, one can either split the clusters and analyze them separately or increase *k* until there is a path between any two points. To achieve the second option one can store the computed distances and increase *k* → *k* + 1. Then any distances that were previously impossible to calculate can be recomputed until no such distances remain.

For all data analysis we chose a small *k* = 4 and used the adaptive version of the algorithm to deal with outliers.

### 4.5 Persistent (co)Homology

*Topological data analysis*, is a rapidly growing field which adapts the computational methods developed for the classification of topological spaces in the late 19th century to the data rich world of today [73, 74, 75]. One of the most popular tools from this field is *persistent (co)homology*, which characterizes the structure in a point cloud by counting the holes that appear and disappear in it on different spatial scales [76, 77].

These computations involve several steps:

1. Given a point cloud *X* embedded in an n-dimensional metric space (ℝ^*n*^, *d*), construct the *Vietoris-Ripps complex V*_*ϵ*_(*X*) by adding an (n-1)-dimensional neighborhood of radius *ϵ* at each point and constructing the following set: *V_ϵ_* = {*xJ_i_*: ∀*J_i_* = {*k,l,m,n*,…}}. *J_i_* is a multi-indexing set whose elements are added if *x_k_, x_i_, x_m_, x_n_* etc. are all a distance *ϵ* from each other. In this context [*x*_*k*_1__, …, *x_k_n__*] denotes an n-simplex and the whole set *V_ϵ_*, corresponds to a *simplicial complex* at a particular choice of scale *ϵ*.
2. Associate the simplices *S_n_* of each dimension to a vector space *C_n_* over a field of coefficients (in this case the field 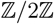 is used).
3. Equip each such vector space with *boundary maps ∂_n_*, with the property that *∂_n_* ₀ *∂_n_* = 0 on any element of any *C_n_*. The collection of these vector spaces along with the corresponding boundary maps is called a chain complex. This chain complex *C*_*_ is visualized with arrows in the following way,

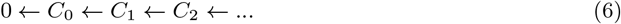
4. If *ϵ* < *δ*, then *V_ϵ_*(*X*) ⸦ *V_δ_*(*X*). This induces an ordering on the generated simplicial complexes and therefore also an ordering on the chain complexes 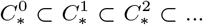 and is called a *persistence complex*. It can be visualized like this,

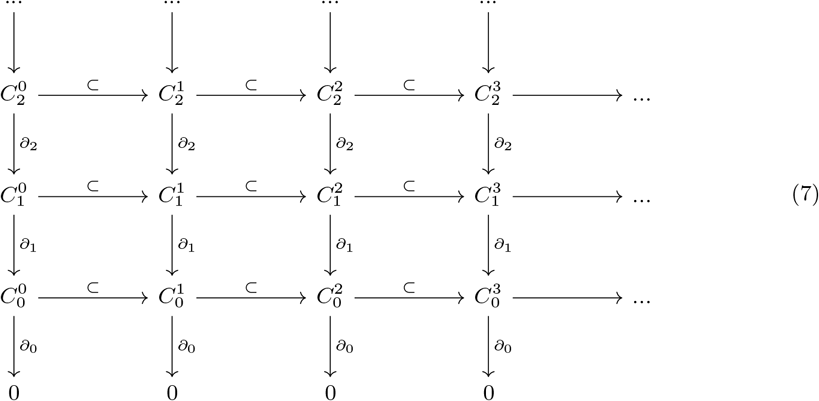 In cohomology the arrows are inverted.
5. Given this construction the next step is to compute the (co)Homology groups associated to the complex at each scale *ϵ*. This is done by computing 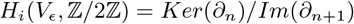. The ranks of the (co)homology groups are known as the Betti numbers, defined by 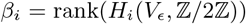, are also computed as a more intuitive summary of the (co)homology groups.
6. Finally, the *ϵ* values at which new (co)homology classes appear are called the births and the values at which they disappear are called deaths. A barcode is the collection of birth-death pairs {(*b_i_,d_i_*)}_*i*_ and large intervals in this collection are interpreted as significant topological features unrelated to noise. From each such interval, the persistence of its corresponding feature is defined as *p_i_* = *d_i_* — *p_i_*

To perform these computations we made use of the openly available Python library *Ripser* [78, 79]. For a more rigorous treatment of algebraic topology see [47, 80].

### 4.6 Statistical testing for Persistent features

While the question of when can a topological feature be considered significant is still an active research area [81, 82], here we stuck to an approach which is highly similar to the ones used in previous applications of topological data analysis to neuroscience. Essentially the idea is that given a point cloud, one can shuffle the responses of each neuron independently and take the feature with maximal persistence **max**{*p_i_*}. By repeating this enough times a distribution of the maximal components is generated, then one can compare the persistent topological features found in real data with the ones occurring in randomly shuffled data.

We only report the results of these statistical tests on one-dimensional topological features, since higher-dimensional features are both unexpected when the input manifold is one-dimensional and we failed to find any such persistent components. For all statistical tests we used 100 shuffles and a Bonferroni corrected p value of 0.0017. Therefore, a topological feature was determined to be significant when it was larger than the 99.83th percentile of the shuffle distribution.

## 5 Acknowledgements

We wish to thank Jonas Verhellen, for his insightful and critical comments during the creation of this manuscript. This research was funded by the European Union’s Horizon 2020 research and innovation programme under the Marie Sklodowska-Curie grant agreement N° 945371 and the University of Oslo.

## 6 Supplementary material

### 6.1 What is a neural manifold?

While the term neural manifold is often used, here we try to provide a slightly more rigorous definition of this concept. First we start with an intuitive definition of a stimulus set.

#### Definition 1

A stimulus set 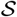 is the set of all stimuli presented to a subject during an experiment. In set notation:

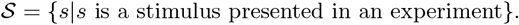

An implicit assumption in most experiments is that for every presented stimulus, highly similar stimuli exist and are “close” to the presented ones in some sense. For example if a pitch of 440Hz is presented there is a highly similar stimulus with a pitch of 440.01Hz. But there is also a highly dissimilar stimulus with a pitch of 20000Hz. This implies that, if one can extend 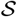 to include these close stimuli, a notion of a metric can be defined which motivates the definition of a stimulus manifold.

#### Definition 2

A stimulus manifold 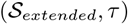 is a topological manifold with the metric topology 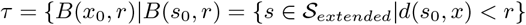 induced by a metric *d*(*x,y*).

Given this definition, we can define the action of a neural population in the following way:

#### Definition 3

A neural population operating on 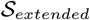, made up of *N* neurons and binned into *T* steps is a mapping 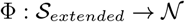, where 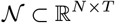.

When transformed by a neural population the closeness relations expressed in the topology of the stimulus manifold *τ* can either be preserved if Φ is a homeomorphism or destroyed, if it is not. Given this fact, one can define a neural manifold as the topological manifold generated by the neural population responses to the stimulus manifold:

#### Definition 4

A neural manifold (**Φ**,*τ*_Φ_) is a topological manifold equipped with the metric topology induced by an appropriate metric *g*(*x, y*).

### 6.2 Supplementary figures

**Figure 5:**
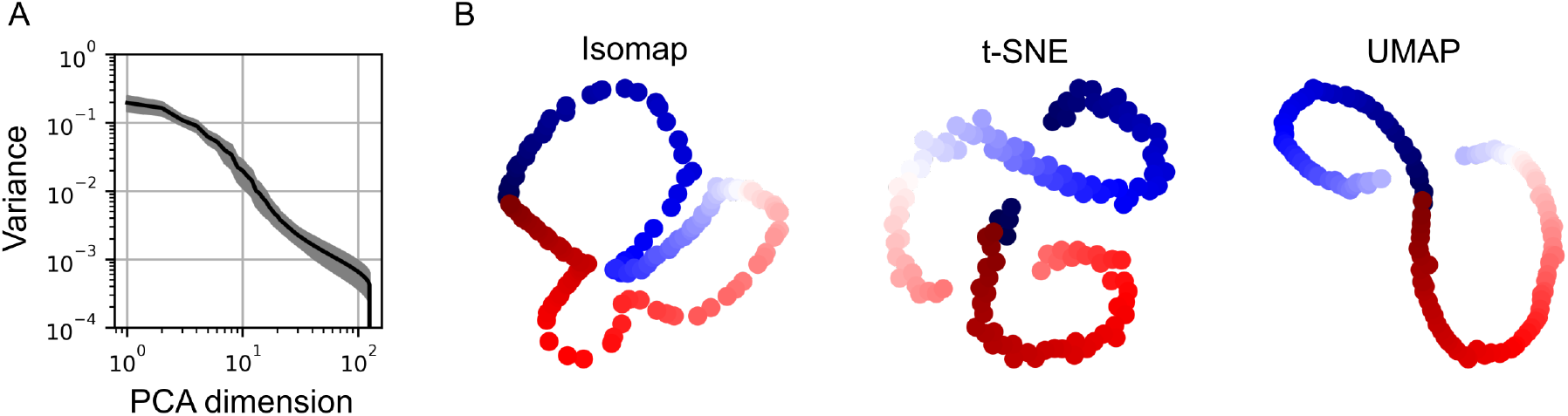
Dimensionality reduction methods. A) Explained variance by principal components averaged over datasets. B) Illustration of manifolds generated by different dimensionality reduction methods, which are distorted from the topologically estimated geometry.

**Figure 6:**
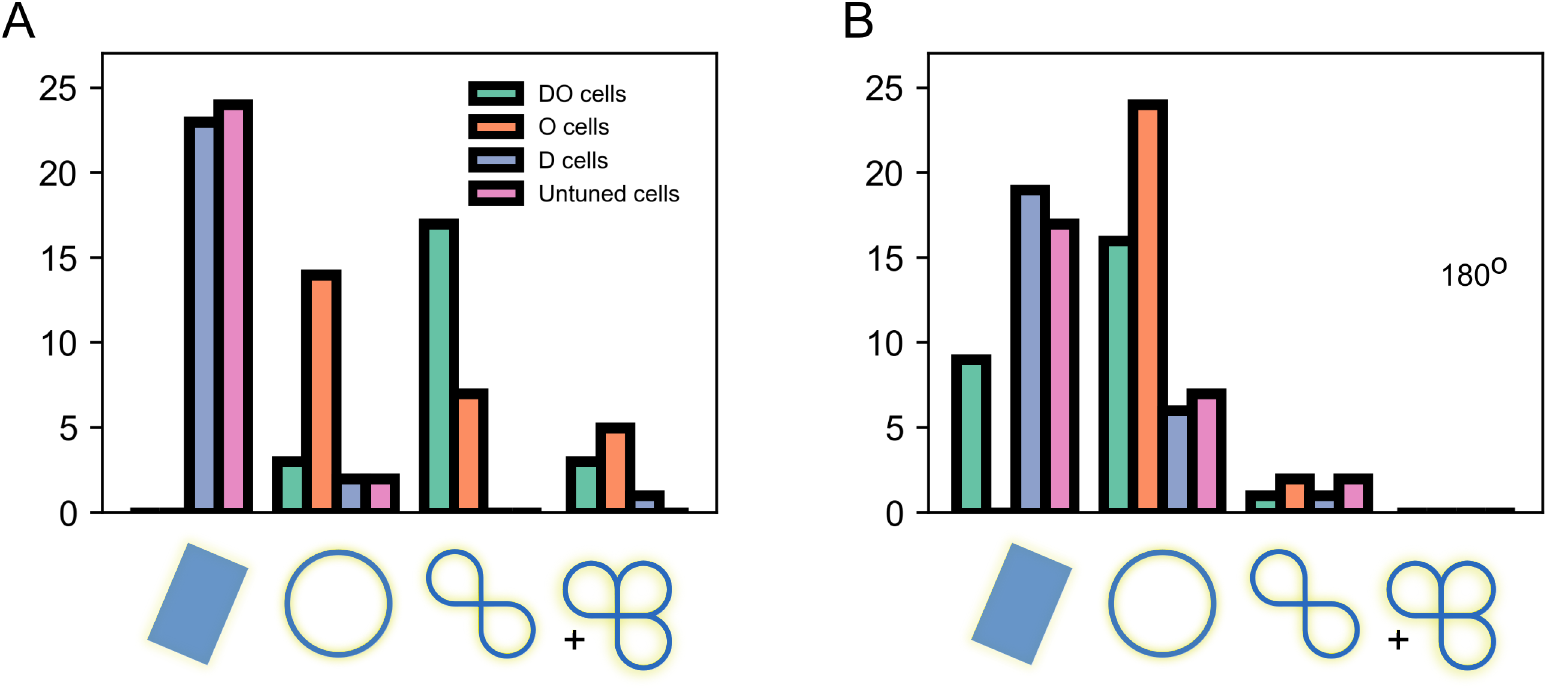
(Co)homological features extracted with the Euclidean metric. A) Histogram of the identified manifolds across all stimuli using the Euclidean metric. While circular features are identified, the high curvature of the neural responses leads to the appearance of too many holes. B) Histograms of the identified holes across subsets of cells. C) Same as B, except the presented angles were limited to 180°. The differences between B and C make the same point as Figure 3, namely O cells encode the rotations on a circle even when the presented angles are restricted. However, the identified topology is much less reliable.

**Figure 7:**
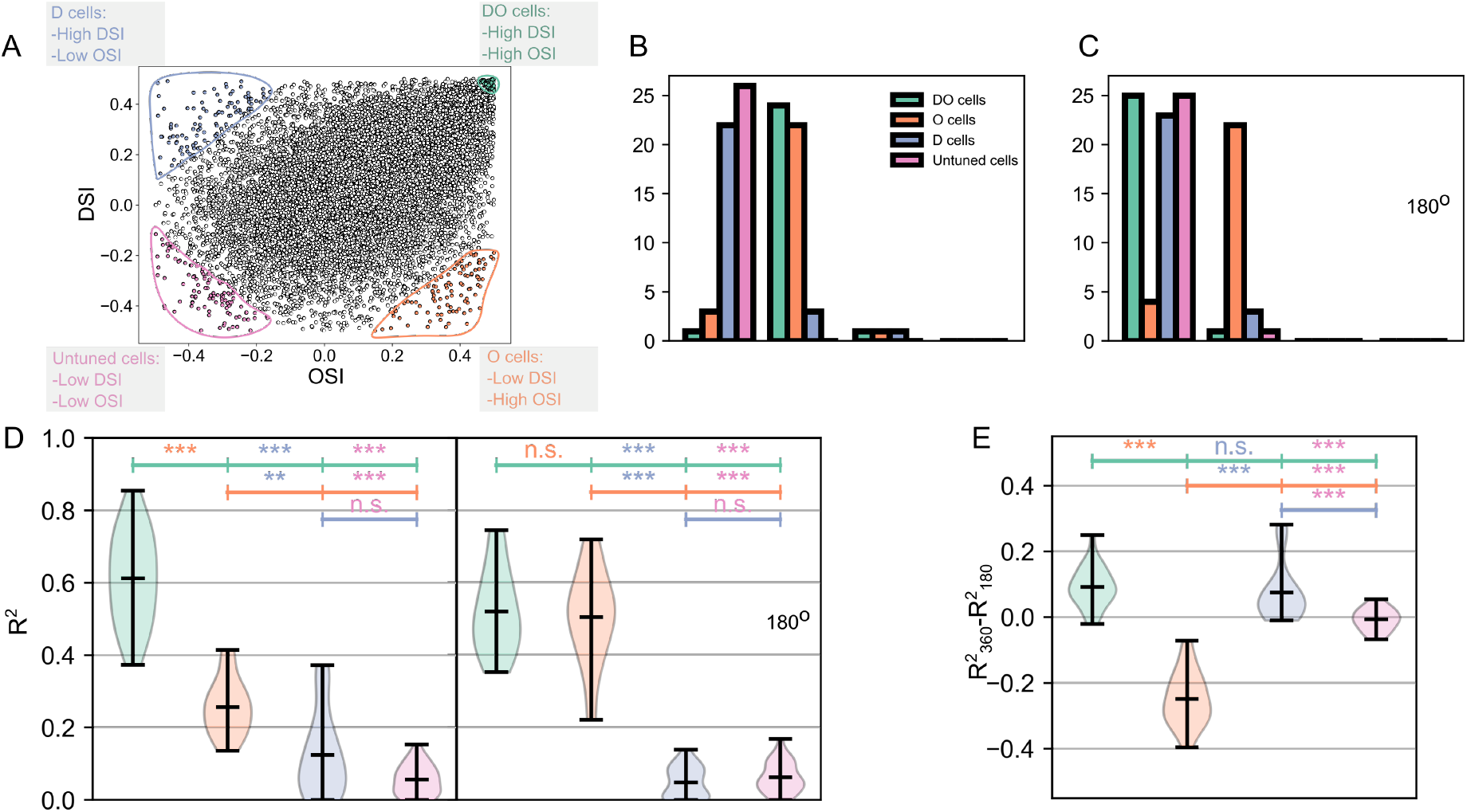
Subpopulations of top 100 strongest tuning cells show similar features. A) Scatter plot showing the identified cell groups. B) Histogram of the identified manifolds for each cell group. C) Same as panel B, except the presented angles were limited to 180° D) Ordinary least squares decoding performance of the different subsets of cells. To determine significance we used a Bonferroni corrected Wilcoxon rank-sum test. *DO* and *O* cells showed good performance, whereas less strongly tuned cells under-performed. The left panel shows predictions for the full 360° of directions, whereas in the right panel the directions were recomputed as *θ* = *θ*mod180°. E) The difference between the predictions showed that only *O* cells exhibit a significant improvement in their performance when it came to recomputed stimuli.

**Figure 8:**
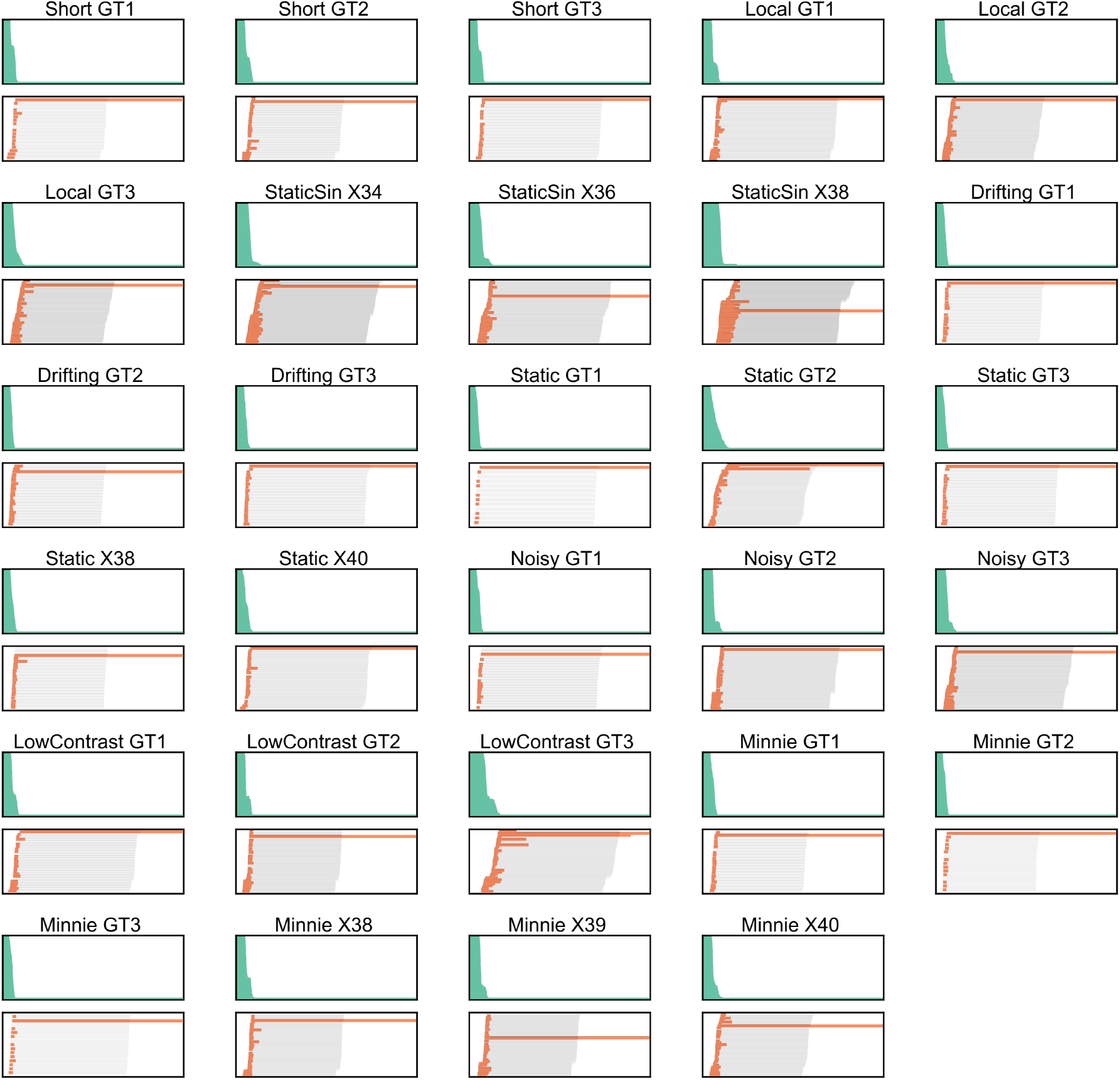
All the barcodes computed with the geodesic metric. The titles denote the stimulus type and the name of the mouse.

